# The lysosomal channel RECS1 regulates extracellular vesicle biogenesis

**DOI:** 10.64898/2026.08.18.745524

**Authors:** Mateus Milani, Nicolas Montes-Bravo, Claudia Sepúlveda-Quiñenao, Jordan Burton, Emiliano Molina, Alvaro Glavic, Birgit Schilling, Lisa M. Ellerby, Claudio Hetz

**Affiliations:** Interdisciplinary Nucleus of Biology and Genetics (NiBG), Institute of Biomedical Sciences, Faculty of Medicine, University of Chile, Santiago, Chile; Biomedical Neuroscience Institute (BNI), Faculty of Medicine, University of Chile, Santiago, Chile; Boldrini Research Center, Campinas, SP, Brazil; Buck Institute for Research on Aging, Novato, CA, 94945, USA; Center for Genome Regulation, Faculty of Sciences, University of Chile, Santiago, Chile

## Abstract

Lysosomal ion channels play key roles in regulating membrane trafficking, autophagy, and cell death. RECS1 is a pH-sensitive lysosomal calcium channel previously implicated in lysosome-mediated apoptosis. Here, we identify a novel role for RECS1 in exosome biology. Using immunoprecipitation followed by mass spectrometry, we mapped RECS1 interactors under apoptotic and lysosomal stress conditions. Notably, Syntenin-1, a key scaffolding protein in ESCRT-independent exosome biogenesis, emerged as the top hit. We validated the physical interaction between RECS1-Syntenin-1 using various approaches. RECS1 localized to secreted exosomes, and its overexpression increased exosome production, as measured by nanoparticle tracking analysis. Intriguingly, a channel-dead RECS1 retained both Syntenin-1 interaction and the ability to promote exosome release, suggesting a channel-independent mechanism. Our findings identify RECS1 as a structural component of the exosomal trafficking machinery and a modulator of extracellular vesicle biogenesis. This work connects lysosomal signalling with intercellular communication and suggests a broader role for RECS1 in stress-responsive secretion.

**TEASER:** RECS1 bridges lysosomal signalling and extracellular vesicles biology, revealing a new mechanism for stress-responsive cell communication.

## INTRODUCTION

Lysosomes have emerged as dynamic signalling hubs that regulate cellular homeostasis and cell death. Lysosomes also function in calcium storage, which can be released through specialized channels to trigger downstream processes such as vesicle fusion, autophagy, and cell death pathway^1,2^ Recently we have identified RECS1 (Responsive to External Centrifugal Stress 1), also known as TMBIM1 (Transmembrane BAX Inhibitor Motif-containing protein 1), as a pH-regulated lysosomal calcium channel^3^. Notably, RECS1 overexpression increases lysosomal acidification and calcium levels, and can induce lysosomal membrane permeabilization (LMP) leading to cell death. Interestingly, RECS1 is the first TMBIM family member shown to promote apoptosis^3^. Given its pro-apoptotic function upon lysosomal damage, we hypothesized that RECS1 might physically interact with apoptotic regulators, such as BAX or other BCL-2 family members, at the lysosome to coordinate cell death signals.

Here we mapped the RECS1 interactome to uncover interacting partners possibly related to cell death pathways. Our unbiased IP-MS interactome screen identified proteins associated with endosomal and exosomal compartments, with Syntenin-1 among the top interactors. Exosomes are small extracellular vesicles of endosomal origin that mediate intercellular communication^4^. Exosome biogenesis is orchestrated by a complex molecular machinery, conducted by two main pathways based on the dependence of ESCRT complexes^5^. In addition to the ESCRT-dependent pathway, the ESCRT-independent pathway mainly involves the adaptor protein Syntenin-1 and its binding partners. Syntenin-1 interacts with the cytosolic tails of syndecans and recruits the accessory protein ALIX facilitating the budding of intraluminal vesicles within multi vesicular bodies (MVBs)^4^. Through the Syndecan–Syntenin–ALIX axis, Syntenin-1 strongly promotes exosome formation and cargo sorting^6^. Importantly, exosomes enriched in Syntenin-1 play significant roles in physio-pathological processes, ranging from developmental signalling to cancer metastasis^5^. In addition, the lysosomes also play an important role in exosome biology, functioning as a main degradation site post-sorting, upon exosome uptake by the receiving cell^7^. Given RECS1 lysosomal localization and the involvement of lysosomes in exosome biology, we considered a possible link between RECS1 and exosome pathways. We therefore characterized RECS1 and Syntenin-1 interaction and asked how RECS1 shapes exosome secretion. Our findings reveal that RECS1 is a cargo of exosomes and also regulates extracellular vesicle biogenesis.

## RESULTS

### IP-MS-based RECS1 interactome identifies exosome biology proteins and highlights Syntenin-1 as the top interactor

To uncover RECS1-associated proteins, we carried out an immunoprecipitation of Flag-tagged RECS1 followed by high-resolution mass spectrometry (Fig. 1A). First, expression of Flag-RECS1 in a HeLa cell Tet-On 3G inducible system was confirmed by immunoblot (Fig. 1B). Because RECS1 expression promotes cell death in cells with lysosomal damage^3^, we performed immunoprecipitation with RECS1-expressing cells treated with lysosomotropic agent chloroquine (CQ) and with wild-type (WT) HeLa cells used as a parental cell control (Fig. 1C). For mass-spectrometry analysis, we compared protein abundance interactions among all the different conditions. Applying a stringent cut-off (log2 fold-change ≥ 3 and q-value ≤ 0.001), we observed that RECS1 pulled down a striking array of endo-lysosomal and exosomal proteins, especially when comparing doxycycline and chloroquine co-treated conditions with other conditions (Tables 1, 2 and 3).

**Figure 1.**
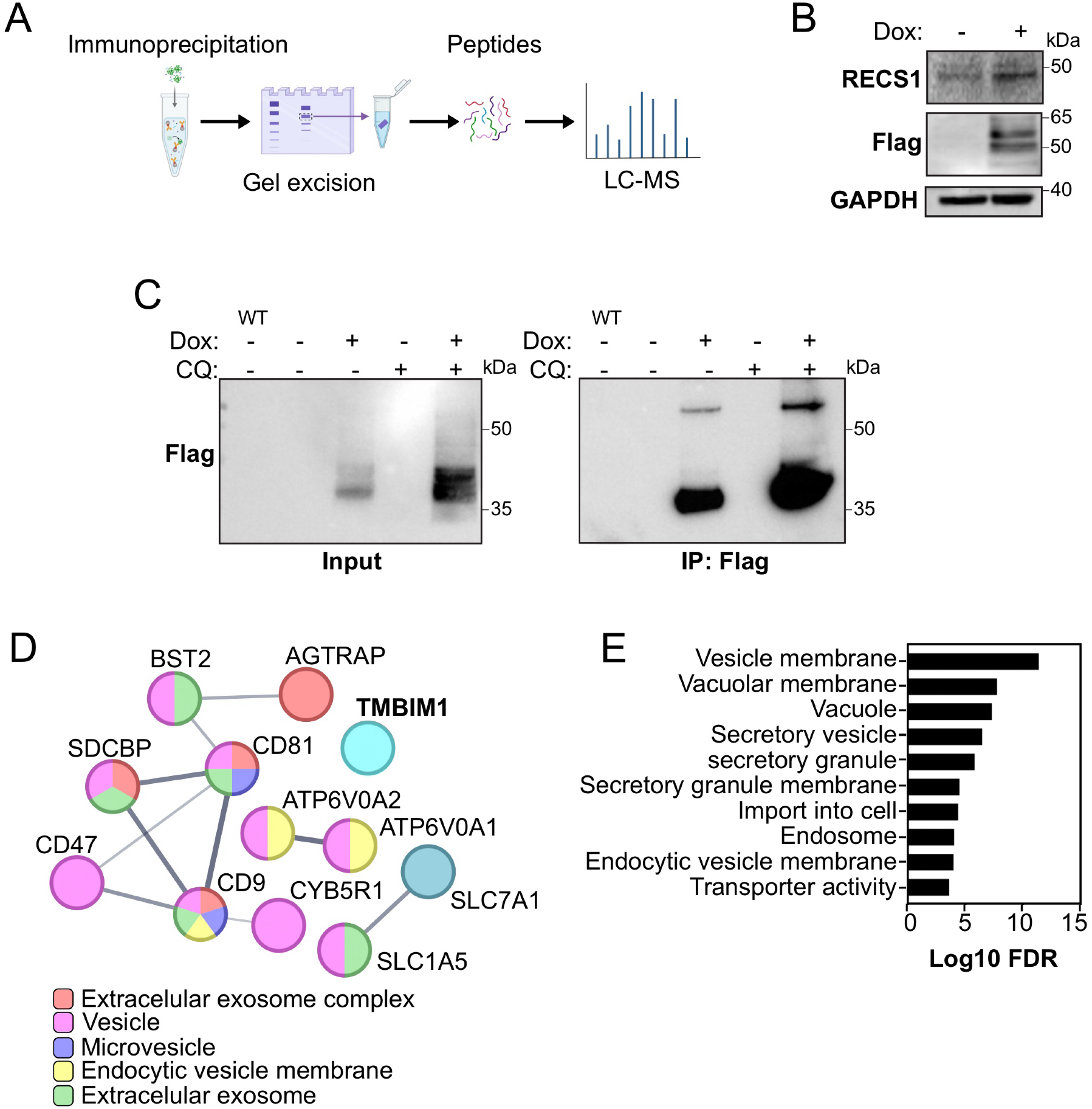
RECS1 interactome reveals exosome biology proteins as main interactors. **(A)** Scheme of IP-MS experimental strategy. **(B)** HeLa Flag-RECS1 cells were stimulated with doxycycline (1 µg/ml, 16 h) and the overexpression of RECS1 and Flag was assessed by immunoblot analysis. **(C)** Flag immunoblot of HeLa WT was used as a control. HeLa Flag-RECS1 cells were stimulated with doxycycline (1 µg/ml, 16 h) and/or chloroquine (100 µM, 24 h) and immunoprecipitated with Flag-bound beads. **(D)** STRING analysis of average log2 ratio of top hits from interactome comparing conditions doxycycline with chloroquine and chloroquine only (k-means clustering = 3); only interconnected nodes, except TMBIM1, are shown. Subcellular localization analyses categories are grouped by color (GO:CC category enrichment). **(E)** Gene Ontology Analysis on the same data as in (D), showing major cellular component categories.

**Table 1.** Average log2 ratio of top hits from interactome between doxycycline with chloroquine and chloroquine only conditions. Exosome-related proteins are highlighted in red.

| Genes | Protein description | AVG.Log2.Ratio |
| --- | --- | --- |
| **TMBIM1** | **Protein lifeguard 3** | **8,63** |
| **SDCBP** | **Syntenin-1** | **4,75** |
| CLCN7 | H(+)/Cl(-) exchange transporter 7 | 4,69 |
| PLP2 | Proteolipid protein 2 | 4,40 |
| ATP6V0A1 | V-type proton ATPase 116 kDa subunit a isoform 1 | 4,39 |
| WBP2 | WW domain-binding protein 2 | 4,10 |
| **CD9** | **CD9 antigen** | **3,87** |
| PPAP2C | Lipid phosphate phosphohydrolase 2 | 3,85 |
| AGTRAP | Type-1 angiotensin II receptor-associated protein | 3,82 |
| M6PR | Cation-dependent mannose-6-phosphate receptor | 3,65 |
| SYPL1 | Synaptophysin-like protein 1 | 3,62 |
| ATP6V0A2 | V-type proton ATPase 116 kDa subunit a isoform 2 | 3,59 |
| TMEM55B | Type 1 phosphatidylinositol 4,5-bisphosphate 4-phosphatase | 3,53 |
| **CD81** | **CD81 antigen** | **3,51** |
| SLC1A5 | Neutral amino acid transporter B(0) | 3,51 |
| BST2 | Bone marrow stromal antigen 2 | 3,47 |
| CYB5R1 | NADH-cytochrome b5 reductase 1 | 3,46 |
| SPPL2A | Signal peptide peptidase-like 2A | 3,41 |
| SLC7A1 | High affinity cationic amino acid transporter 1 | 3,35 |
| **CD47** | **Leukocyte surface antigen CD47** | **3,23** |
| DAGLB | Sn1-specific diacylglycerol lipase beta | 3,14 |
| PIEZO1 | Piezo-type mechanosensitive ion channel component 1 | 3,14 |
| SYNGR2 | Synaptogyrin-2 | 3,13 |
| SLC5A6 | Sodium-dependent multivitamin transporter | 3,07 |

**Table 2.**
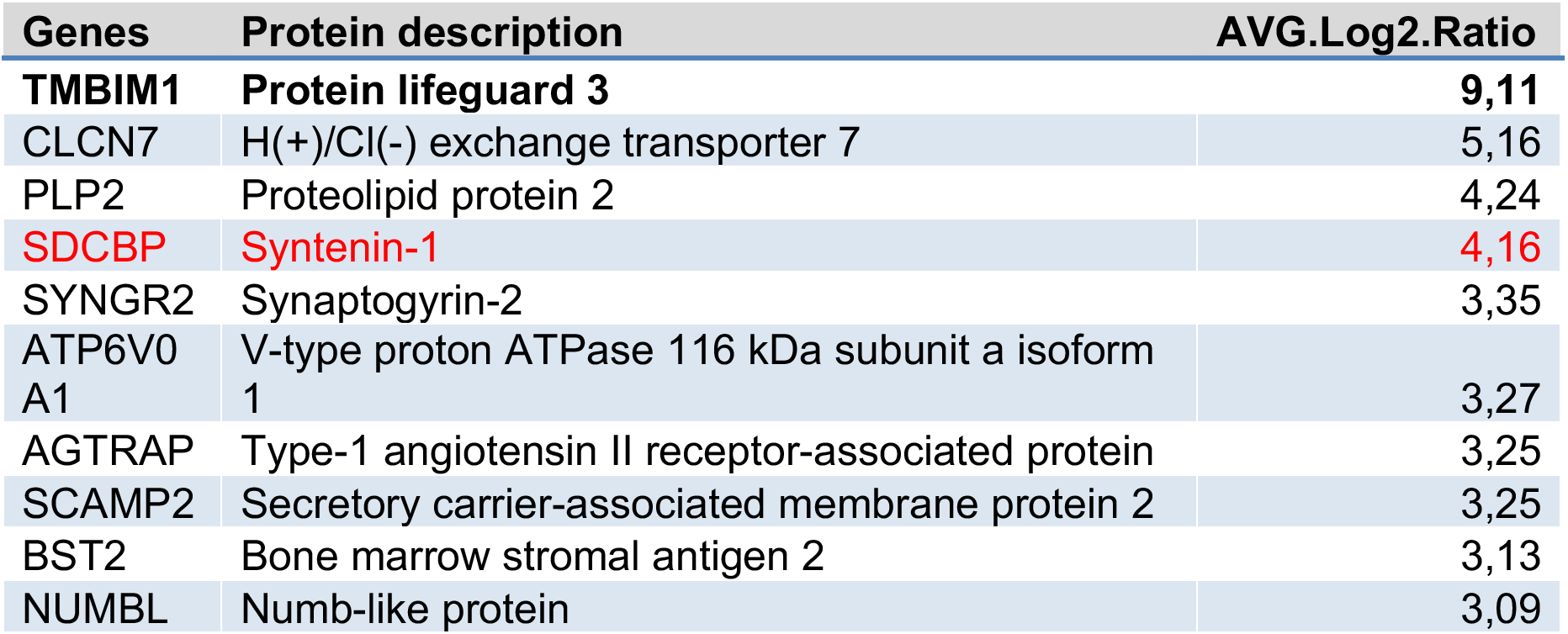
Average log2 ratio of top hits from interactome between doxycycline and vehicle conditions. Exosome-related proteins are highlighted in red.

| Genes | Protein description | AVG.Log2.Ratio |
| --- | --- | --- |
| **TMBIM1** | **Protein lifeguard 3** | **9,11** |
| CLCN7 | H(+)/Cl(-) exchange transporter 7 | 5,16 |
| PLP2 | Proteolipid protein 2 | 4,24 |
| **SDCBP** | **Syntenin-1** | **4,16** |
| SYNGR2 | Synaptogyrin-2 | 3,35 |
| ATP6V0<br>A1 | V-type proton ATPase 116 kDa subunit a isoform<br>1 | 3,27 |
| AGTRAP | Type-1 angiotensin II receptor-associated protein | 3,25 |
| SCAMP2 | Secretory carrier-associated membrane protein 2 | 3,25 |
| BST2 | Bone marrow stromal antigen 2 | 3,13 |
| NUMBL | Numb-like protein | 3,09 |

**Table 3.**
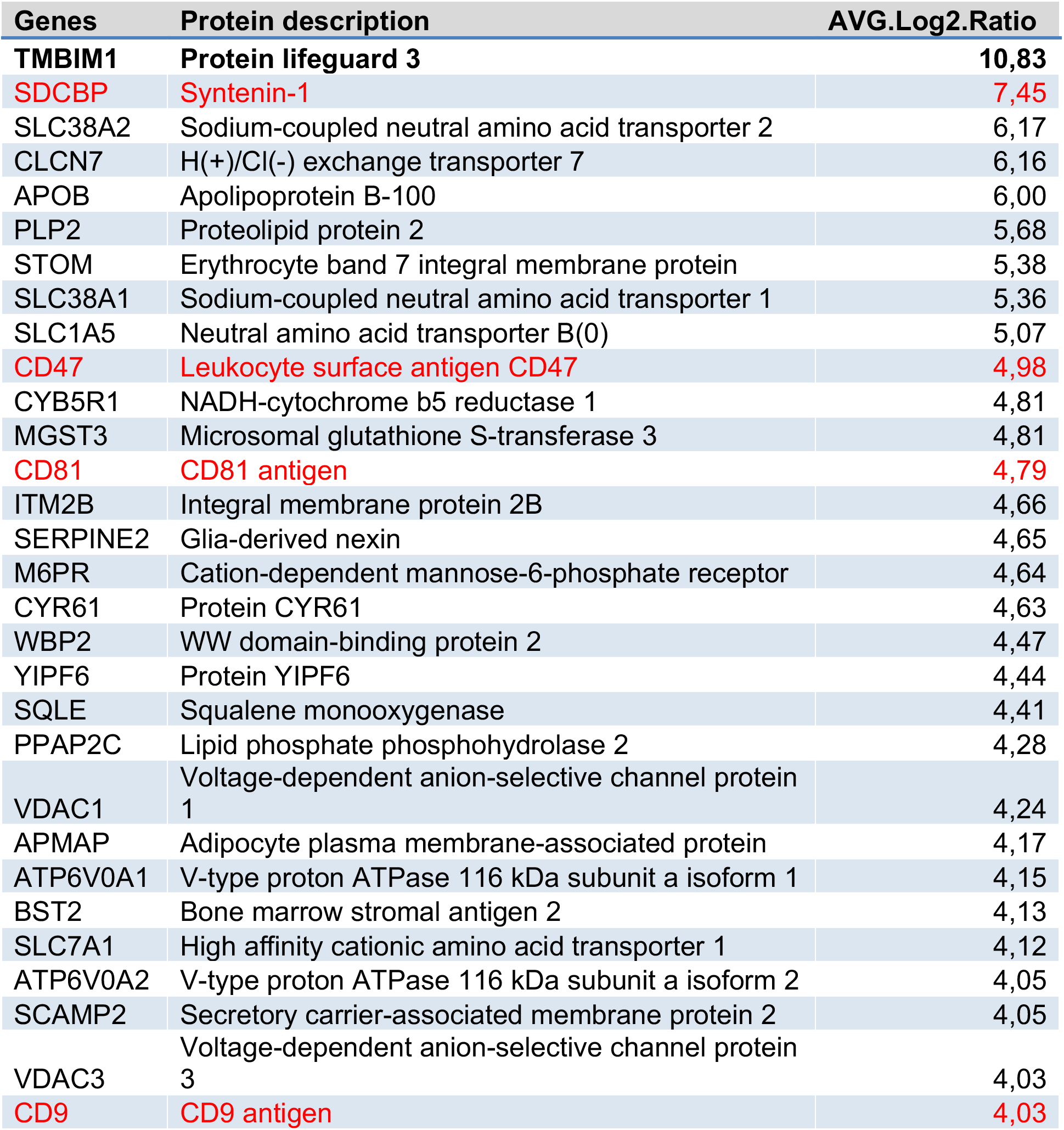
Average log2 ratio of top hits from interactome between doxycycline with chloroquine and vehicle conditions. Exosome-related proteins are highlighted in red.

Surprisingly, no BCL-2 family proteins or classic apoptosis regulators were significantly enriched, indicating that RECS1 does not stably bind apoptotic factors under these conditions. Instead, multiple exosome-related proteins were highly enriched in the RECS1 interactome. Of those, Syntenin-1 was the most prominent interactor after the bait itself. Moreover, other known components of exosome biogenesis and regulators of cargo sorting were also pulled down with RECS1. These include tetraspanins CD9 and CD81, classical exosomal membrane markers^8^; the endosomal H^+^/Cl^−^ exchanger CLCN7^9^ and V-ATPase subunits ATP6V0A1/A2^10^, involved in endolysosomal acidification^11^; and membrane trafficking proteins such as PLP2^12^ and SCAMP2^13^. STRING network analysis of the high-confidence hits showed a tightly interconnected cluster of proteins with functions in exosome and vesicles categories (Fig. 1D). Gene Ontology enrichment confirmed that extracellular vesicle components were overrepresented among RECS1 partners (Fig. 1E). Together, these results reveal that RECS1 is associated with the molecular machinery of exosome biogenesis, with Syntenin-1 emerging as a central candidate, suggesting that RECS1 might interact with the Syntenin-1–related exosomal machinery.

### RECS1 is present at exosomal vesicles and co-localises with Syntenin-1 at endolysosomal compartments

Our interactome contained several proteins associated with multivesicular bodies (MVB) formation and exosome biogenesis, raising the possibility that a fraction of RECS1 resides on intraluminal vesicles (ILVs) destined for secretion. To test this, we asked whether RECS1 is detected in secreted small EVs and whether it localizes to late endosomes/MVBs. Extracellular vesicles were purified from conditioned cell culture medium of control and RECS1-expressing HeLa cells using an established multi-step protocol^14^ that allowed us to isolate microvesicles and exosomes (Fig. 2A). Western blot analysis of these fractions demonstrated that overexpressed RECS1 is highly present in the exosome fraction, along with Syntenin-1 (Fig. 2B). Conversely, the exclusion marker Calnexin^15^ was present in cell lysates but not in exosomal fraction, confirming the purity of the assay. These data indicate that a portion of RECS1 resides on secreted exosomes, which is consistent with the interactome revealing RECS1 interaction with exosomal proteins.

**Figure 2.**
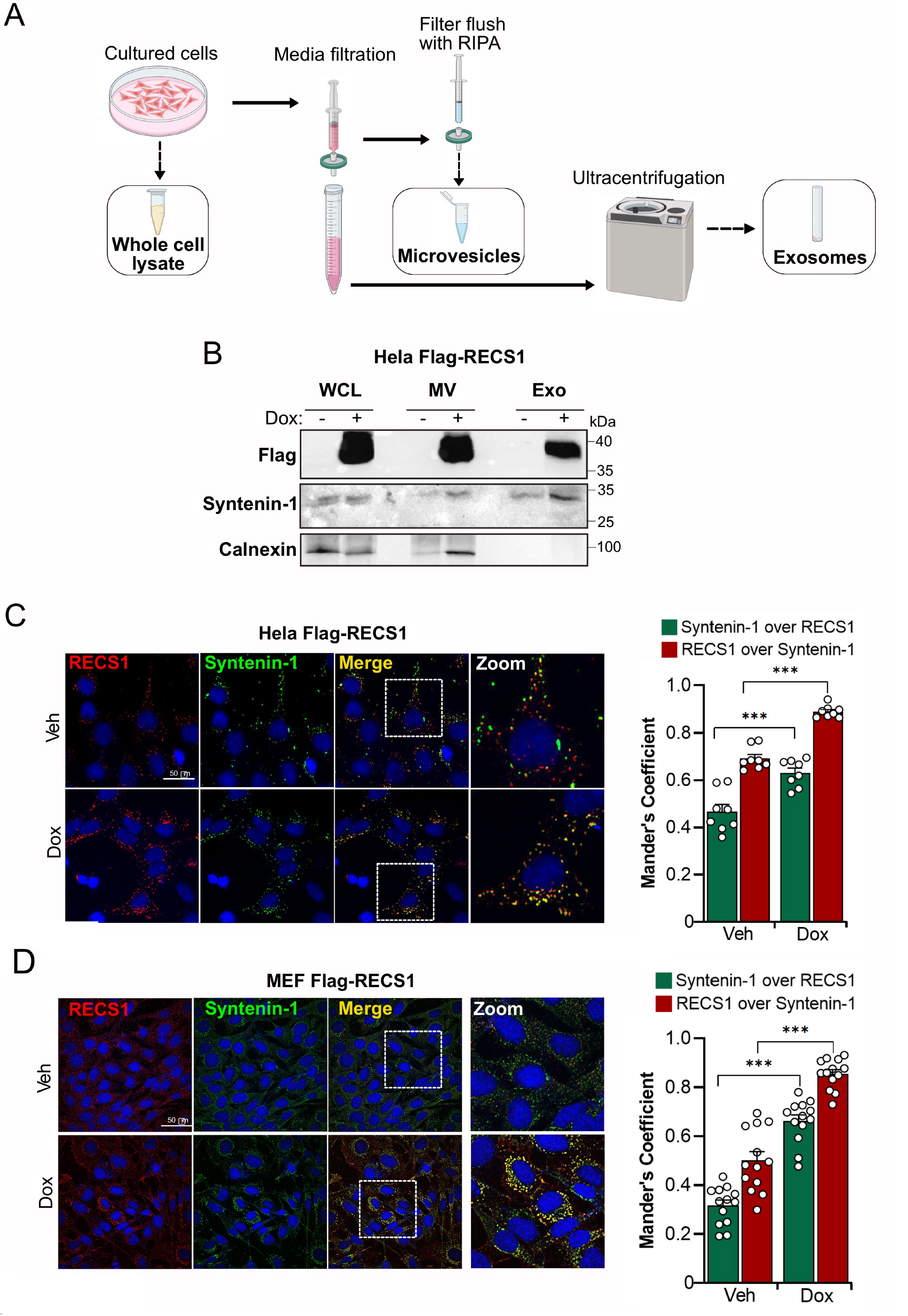
RECS1 is located at exosomal vesicles and co-localises with Syntenin-1. **(A)** Scheme representing the vesicle isolation protocol. **(B)** HeLa Flag-RECS1 were stimulated with doxycycline (1 µg/ml, 16 h), processed for extracellular vesicles and exosome purification, and indicated proteins were assessed by immunoblot. WCL: whole cell lysate; MV: microvesicles; Exo: exosomes. **(C)** HeLa Flag-RECS1 and **(D)** MEF Flag-RECS1 were stimulated with doxycycline (1 µg/ml, 16 h), processed for immunocytochemistry and stained for RECS1 and Syntenin-1. <u>Left</u>: representative confocal images. <u>Right</u>: co-localization quantification by Mander’s coefficient; green depicts the percentage of signal of Syntenin-1 over RECS1 and red depicts the percentage of signal of RECS1 over Syntenin-1.

Given that Syntenin-1 was the top exosomal interactor of RECS1 in our IP-MS, we examined the intracellular distribution of these two proteins. Confocal immunofluorescence microscopy revealed a substantial co-localization of RECS1 with Syntenin-1 in punctate vesicular structures consistent with endosomes or lysosomes compartments in both HeLa and MEF cells (Fig. 2C and D). In cells overexpressing RECS1, a predominantly perinuclear pattern that overlapped with Syntenin-1 was revealed. Quantitative analysis demonstrated a high degree of co-localization with a large fraction of RECS1-positive vesicles containing Syntenin-1, and vice versa (Fig. 2C and D, right panels. Altogether, these results suggest that RECS1 is present at secreted exosomes and that both RECS1 and Syntenin-1 are present on endolysosomal compartments.

### RECS1 promotes exosome secretion and physically interacts with Syntenin-1

Next, we asked if RECS1 affects the amount of exosomes released by cells. Quantification of purified exosomes by Nanosight revealed a significant increase in particle concentration released by RECS1-overexpressing cells (Fig. 3A), which was further confirmed upon normalisation to protein content on the whole cell lysate from which the exosomes were derived (Fig. 3B). These results suggest that RECS1 actively enhances the biogenesis and/or secretion of exosomes.

**Figure 3.**
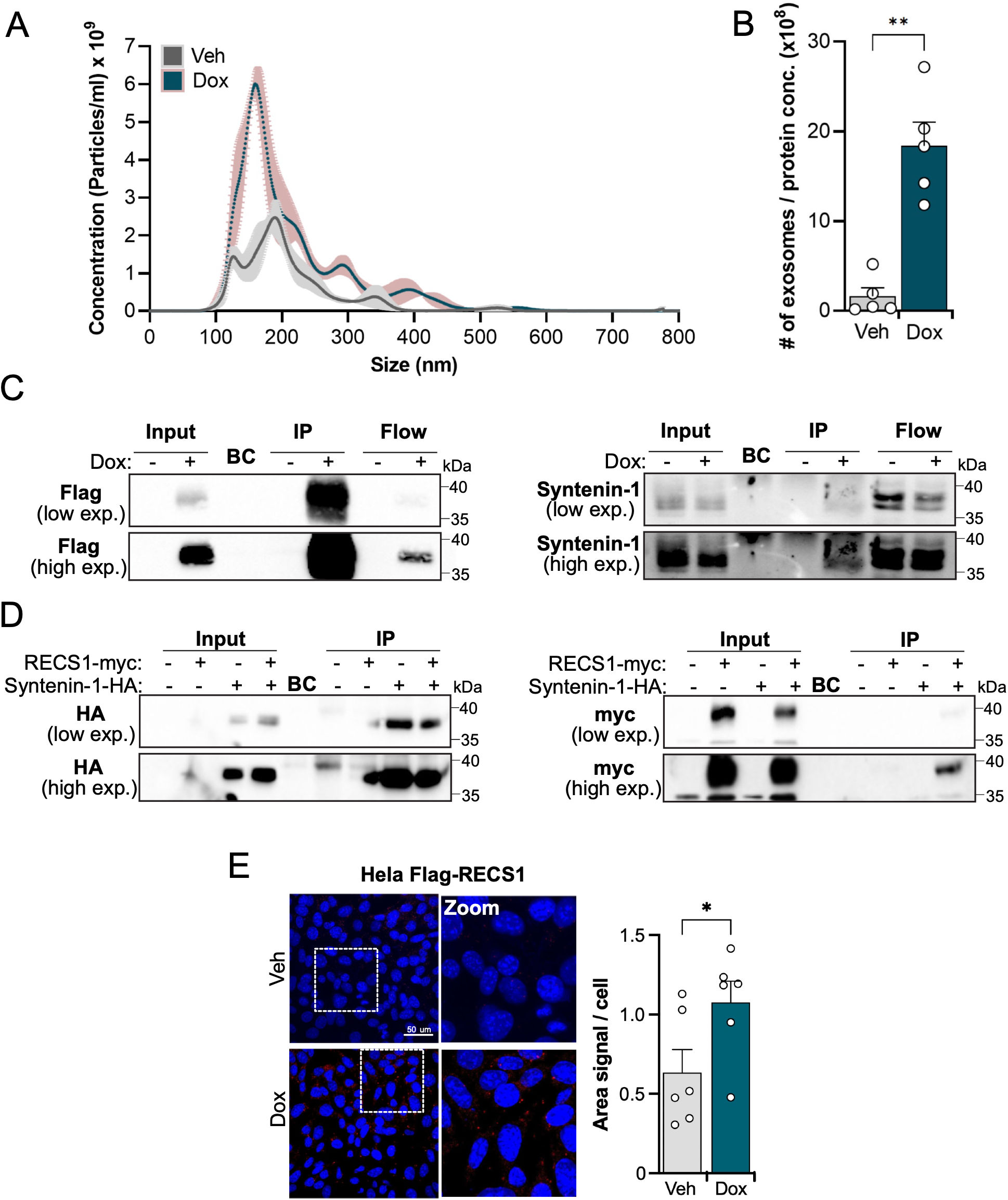
RECS1 promotes exosome secretion and interacts with Syntenin-1. **(A)** Representative Nanosight plot of purified exosomes from HeLa Flag-RECS1 stimulated with doxycycline (1 µg/ml, 16 h) **(B)** The number of exosomes were normalized to WCL protein concentrations (n = 5). **(C)** HeLa Flag-RECS1 cells were stimulated with doxycycline (1 µg/ml, 16 h) and processed for Flag immunoprecipitation. **(D)** HEK-293 cells transfected with RECS1-myc and/or Syntenin-1-HA plasmids were processed for HA immunoprecipitation. **(E)** HeLa Flag-RECS1 were stimulated with doxycycline (1 µg/ml, 16 h) and subjected to proximity ligation assay. <u>Right</u>: representative confocal images. <u>Left</u>: quantification of the ratio from area of PLA signal to the total number of cells.

To determine if RECS1 and Syntenin-1 are part of a protein complex, we performed co-immunoprecipitation (co-IP) experiments. In HeLa cells expressing Flag-RECS1, pulling down RECS1 efficiently co-precipitated endogenous Syntenin-1 (Fig. 3C). Specificity was confirmed by detecting Syntenin-1 in doxycycline-induced Flag-RECS1 IP samples, but not in control IPs from non-induced cells. Furthermore, in a transient transfection system, we co-expressed RECS1-myc and Syntenin-1-HA in HEK293 cells and immunoprecipitated Syntenin-1 via its HA tag. We found that RECS1 co-precipitated with Syntenin-1 in this heterologous system as well (Fig. 3D), as evidenced by the presence of RECS1-myc in anti-HA pull-downs only when both proteins were co-expressed. Similar interactions could be detected with the myc-based immunoprecipitation (Supplementary Fig. S1). These co-IP results validate that RECS1 and Syntenin-1 associate biochemically, either through a direct interaction or as part of a larger protein complex.

The RECS1- Syntenin-1 protein interaction was also confirmed using a proximity ligation assay. Robust fluorescent PLA puncta were observed in cells with RECS1 expression, indicating close proximity of RECS1 and Syntenin-1 molecules in HeLa cells (Fig. 3E). Altogether, these data provide evidence that RECS1 and Syntenin-1 interact intracellularly, supporting the idea that RECS1 engages with the exosomal biology machinery via Syntenin-1.

### RECS1 channel activity is dispensable for exosome production and Syntenin-1 interaction

Previously, we have shown that the RECS1 D295Q mutation, which abolishes its channel function, fully ablated its pro-apoptotic function under lysosomal stress^3^. To determine whether RECS1 ion channel activity is required for its association with exosome biology, we used an inducible cell line that conditionally expresses Flag-RECS1 D295Q. First, we confirmed the mutant phenotype by comparing its ability to prevent apoptosis with WT RECS1 expressing cells. As reported, overexpression of the D295Q mutant showed reduced cell death either alone by overexpression or after chloroquine treatment (Supplementary Fig. S2A).

We asked if the presence of RECS1 in the exosomal fractions were dependent on its channel function. Interestingly, RECS1 was found in great amounts in the exosomal fractions of Flag-RECS1 D295Q cells (Fig. 4A), suggesting RECS1 involvement in exosome biology is independent of its channel properties. Next, similarly to our previous experiments with the functional RECS1, we quantified exosome production in cells expressing RECS1 D295Q. Again, the channel-dead mutant did not impair exosome production, reflected in a similar increase in exosome production when RECS1 D295Q was expressed when compared with WT RECS1 (Figs. 4B and C).

**Figure 4.**
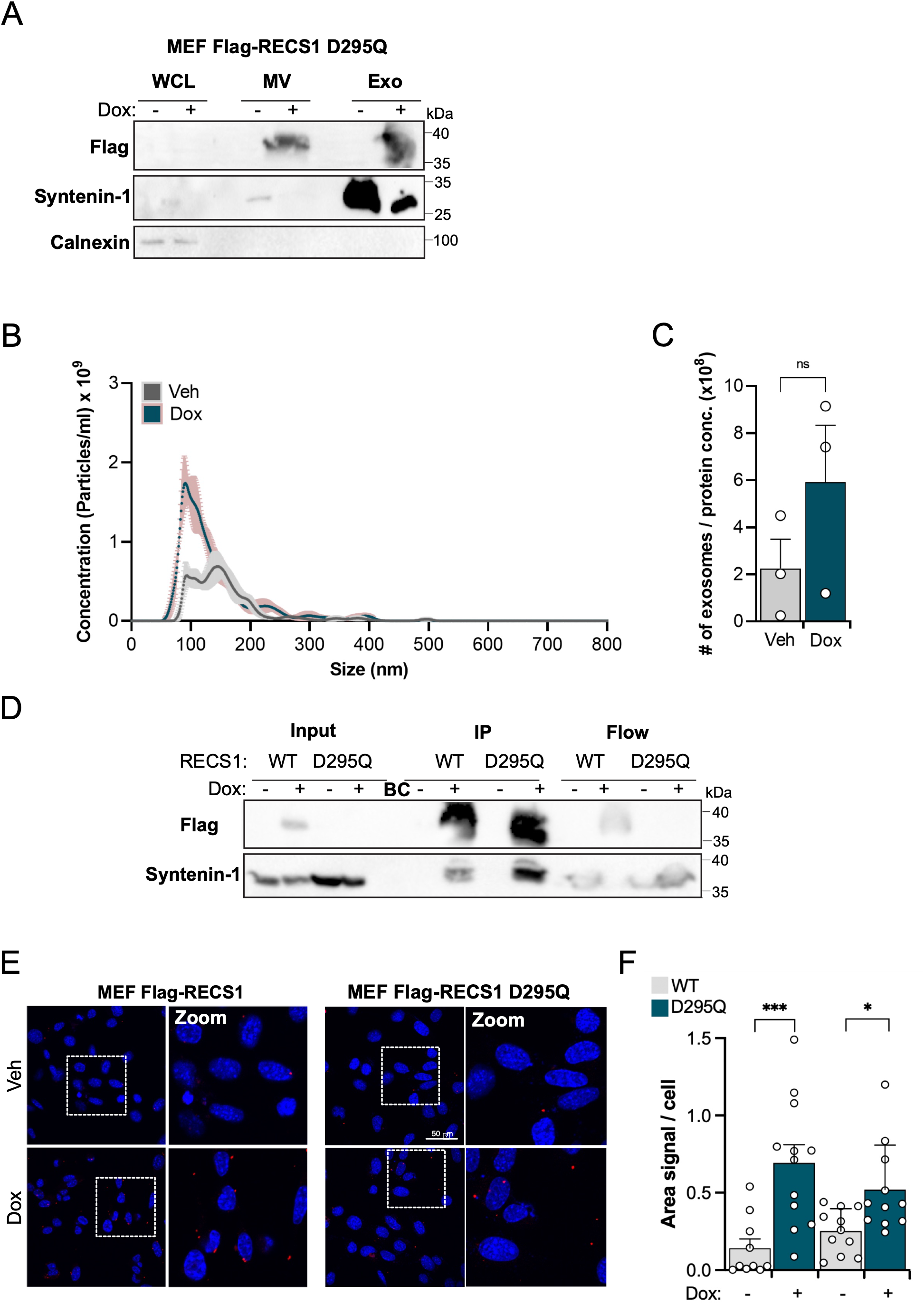
RECS1 channel activity is not required for exosome production nor interaction with Syntenin-1. **(A)** MEF Flag-RECS1 D295Q were stimulated with doxycycline (1 µg/ml, 16 h), processed for extracellular vesicles and exosome purification and indicated proteins were assessed by immunoblot. WCL: whole cell lysate; MV: microvesicles; Exo: exosomes. **(B)** Representative Nanosight plot of purified exosomes from MEF Flag-RECS1 D295Q stimulated with doxycycline (1 µg/ml, 16 h). **(C)** The number of exosomes were normalized to WCL protein concentrations (n = 3). **(D)** MEF Flag-RECS1 WT and D295Q cells were stimulated with doxycycline (1 µg/ml, 16 h) and processed for Flag immunoprecipitation. **(E)** Representative confocal images of MEF Flag-RECS1 WT and D295Q stimulated with doxycycline (1 µg/ml, 16 h) and subjected to proximity ligation assay. **(F)** Quantification of the ratio from area of PLA signal to the total number of cells of MEF depicted in E.

We next examined whether the D295Q mutation affected RECS1 ability to interact with Syntenin-1. Although immunocytochemistry co-localization analysis indicated that RECS1 D295Q signal does not overlap with Syntenin-1 as much as with WT RECS1 (Supplementary Fig. S2B), side-by-side comparison by immunoprecipitation of WT and D295Q RECS1 clearly show interaction between RECS1 and Syntenin-1 in both cells upon doxycycline exposure (Fig. 4D). These results were further confirmed by side-by-side PLA, which showed a strong signal indicating close proximity between RECS1 and Syntenin-1 in both WT and D295Q mutant (Fig. 4E). Altogether, these results suggest that RECS1 interaction with Syntenin-1 and its role in promoting exosome secretion do not require its channel activity, pointing instead to a structural or scaffolding role for RECS1 under these conditions.

### RECS1 increases exosome production in vivo

Since RECS1 is involved in the formation of MVBs in humans through the recruitment of components of the ESCRT machinery^16,17^, which is required for MVB formation^18–20^, and our data suggested an involvement of RECS1 in exosomal biology, we moved forward to evaluate the evolutionary conservation of RECS1 on this process using *D. melanogaster* as a model (dRecs1). We performed time-and tissue-specific gain-and loss-of-function of dRecs1 using the Gal4/UAS tool^21^, using an adult tissue called accessory gland (AG) as a model, to evaluate its role in MVB formation (Fig. 5A). This tissue is related to the mammalian prostate, with a sac-like structure which formed by an epithelium comprised of 2 cell types: main cells (MCs) and secondary cells (SCs). The latter are characterized by a large development of MVBs and have been shown to generate the exosomes/extracellular vesicles (EVs) that are secreted into the lumen of the gland^22,23^. In *D. melanogaster*, these EVs bind to both sperm and epithelial cells of the female tract after intercourse. These vesicles carry, among other things, peptides that alter the behavior of females, preventing remating behavior^23–25^.

**Figure 5.**
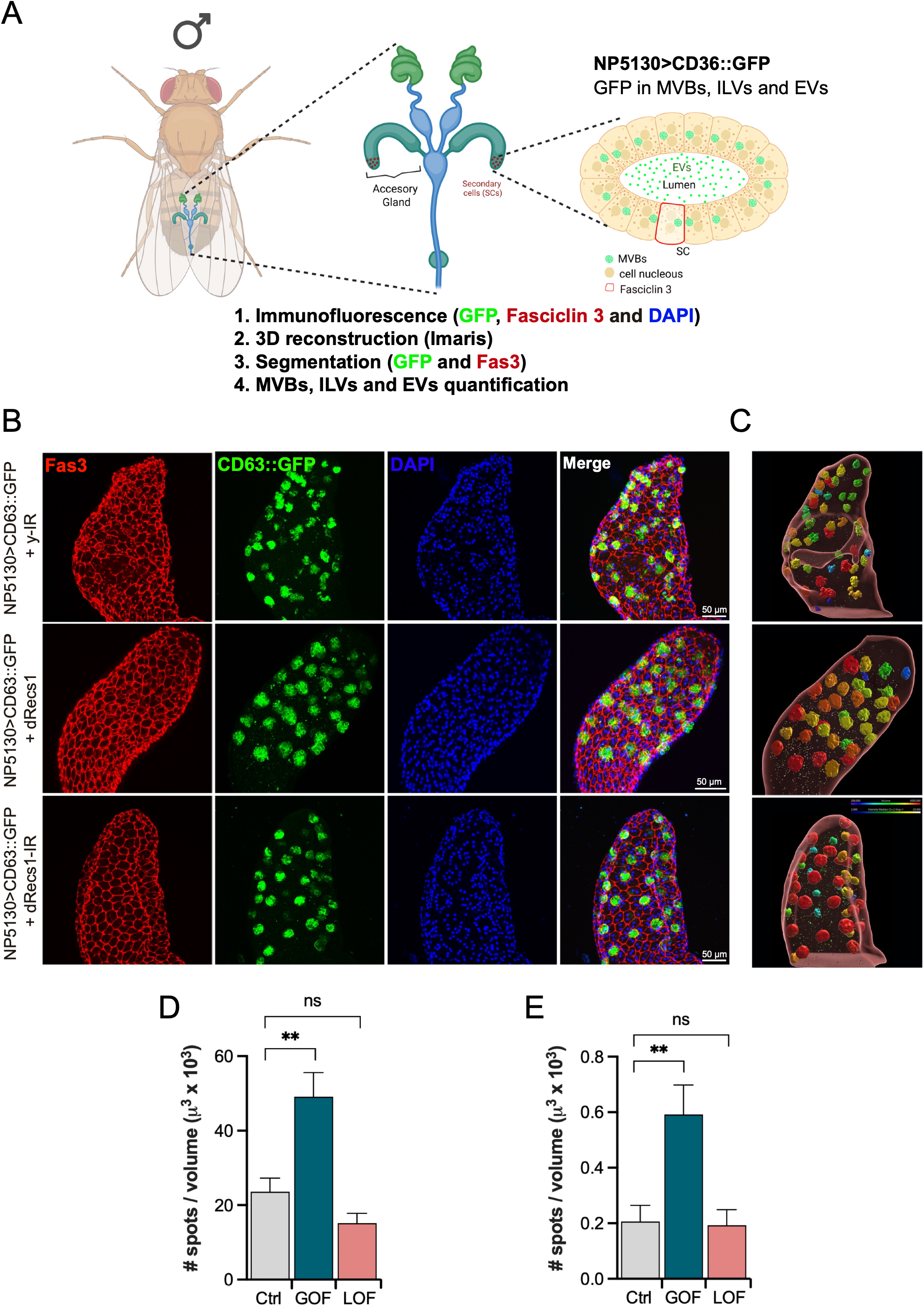
RECS1 overexpression induces exosomal production in *D. melanogaster*. **(A)** Scheme of methodology used to analyze accessory glands of *D. melanogaster* males. Gal4/UAS system was used to direct expression in AGs (NP5130 driver). After the dissection of AGs, immunofluorescence was performed. Septate junctions at the cell surface were stained with Fasciclin 3 (Fas3), CD63 fused to GFP is present in intraluminal vesicles, multivesicular bodies (MVBs) and extracellular vesicles (EVs) (CD63::GFP). Finaly nuclei were stained with DAPI. 3D renderings were performed to segment the accessory glands and quantify the EVs secreted into the lumen. **(B)** Immunofluorescences of accessory glands co-expressing CD63 fused to EGFP (CD63::EGFP) with: RNA interference against yellow (y-IR; Control), dRecs1 fused to Flag epitope (dRecs1, GOF) and RNA interference against dRecs1 (dRecs1-IR; LOF). Fasciclin 3 (Fas3, red), CD63 fused to GFP (CD63::GFP, green) and nuclei (DAPI, blue). A representative image is presented for each condition. Scale bar= 50 µm. (**C)** 3D renderings of representative AGS for each condition were generated to visualize the gland surface, MVBs and EVs spots. **(D)** Quantification of ILVs and MVBs number normalized by segmentation volume. **(E)** Quantification of EVs number normalized by segmentation volume.

To evaluate the role of dRecs1 in MVB formation and EV release, we expressed an interference RNA against dRecs1 (dRecs1-IR) to assess loss of function and overexpression of the dRecs1-Flag fusion protein as a gain-of-function strategy. Both conditions were co-expressed with the CD63::GFP fusion protein, which labels both MVBs and EVs^24,26^, followed by quantification of MVBs and EVs in immunofluorescence samples from adult accessory glands (Fig. 5A). Interestingly, overexpression of dRecs1 induces a significant increase in the number of EVs in the lumen of the accessory gland, in line with our results *in vitro*. On the other hand, knockdown of Recs1 (dRecs1-IR) did not show any evident alteration in the number of EVs compared to the control condition (expression of an RNA interference against the yellow gene, y-IR). Moreover, the increase in the number of EVs correlated with an increase in the intracellular CD63::GFP signal (intraluminal vesicles and MVBs), suggesting that MVB production may also be promoted by dRecs1 (Figure 1B-E). These observations are consistent with a previous report that hRecs1 promotes MVB formation through its physical interaction with tumor susceptibility 101 (TSG101), a subunit of the ESCRT-I complex^16,27^. Notably, dRecs1 lacks the PSAP motif, suggesting that its ability to stimulate MVB formation may not solely depend on RECS1 interaction with the ESCRT machinery through this motif. Altogether, our *in vivo* data support our results in vitro, highlighting a novel function for RECS1 in the control of exosomal biogenesis.

## DISCUSSION

In this study, we describe a novel role for the lysosomal protein RECS1 as a modulator of exosome biology. Through an unbiased interactome analysis, we identified that RECS1 physically associates with multiple proteins involved in exosome biogenesis and trafficking, with Syntenin-1 emerging as its most prominent interactor. We validated this protein-protein interaction through co-immunoprecipitation, immunofluorescence-based colocalization, and proximity ligation assays. Functional assays further revealed that RECS1 localizes to exosomal fractions and enhances exosome secretion. These findings represent a significant shift in our understanding of RECS1 role in cell biology beyond its known involvement in lysosome-mediated cell death and highlight its participation in intercellular communication through extracellular vesicle release.

The identification of Syntenin-1 as a RECS1 interactor provided a compelling lead into exosome biology. Syntenin-1 plays a central role in ESCRT-independent exosome biogenesis, linking syndecans to ALIX and promoting the inward budding of intraluminal vesicles within multivesicular bodies^6,16^. Importantly, we demonstrated that overexpression of RECS1 increases exosome output and that RECS1 is present within purified exosomal fractions, along with canonical EV markers. These results support the hypothesis that RECS1 actively participates in exosome production.

Intriguingly, the RECS1 D295Q mutant, which lacks its ion channel activity, retained the ability to interact with Syntenin-1 and to promote exosome release. This result challenges the assumption that RECS1 function in vesicle biology is mediated by its channel properties. Instead, it suggests that RECS1 may play a structural or scaffolding role in exosome biogenesis. Although RECS1 is recognized as a pH-sensitive calcium channel, its association with the exosomal machinery seems to be independent on calcium flux or ion permeability by RECS1. These findings distinguish RECS1 from other lysosomal channels such as TRPML1, which role in exosome secretion is directly linked to calcium signaling^17,18^. One possible explanation for this is that RECS1 may serves as a membrane platform that facilitates the assembly of protein complexes involved in vesicle budding or cargo sorting. The interaction with Syntenin-1 may be direct, potentially involving a PDZ-like-binding motif on RECS1 cytosolic tail, although this remains to be experimentally validated. Alternatively, RECS1 might influence the curvature or lipid composition of endosomal membranes in ways that favor intraluminal vesicle formation, which could be occurring by altering local membrane tension or charge. In this context, the recruitment of Syntenin-1 to specific membrane subdomains could be stabilized or enhanced by RECS1 presence, thereby modulating the assembly of the Syntenin–ALIX complex.

It has been shown that when lysosomal degradation is impaired, cells increase exosome release as a compensatory mechanism to offload undigested material^19,20^. In this context, RECS1 may serve as a lysosomal sensor or modulator that facilitates this trafficking switch. Indeed, our interactome comparisons show that the presence of chloroquine enhances the enrichment of exosomal proteins in RECS1 immunoprecipitations, suggesting that stress conditions potentiate RECS1 engagement with exosome-related pathways.

Functionally, the involvement of RECS1 in exosome production could have significant implications in both physiological and pathological processes. Exosomes play key roles in cancer progression, immune modulation, and neurodegenerative diseases^21–23^. For example, in the tumor microenvironment, exosomes can disseminate oncogenic signals, alter stromal responses, and contribute to metastasis^24^. Syntenin-1 itself has been implicated in several cancers as a driver of exosome-mediated communication^25^. It is plausible that RECS1, through its regulation of exosome release, could influence such processes. Conversely, in the context of neurodegeneration, exosome release has been proposed as a mechanism for clearing aggregation-prone proteins such as α-synuclein and tau^26,27^. In this context, if RECS1 facilitates this form of lysosome-to-exosome trafficking, it might play a protective role under conditions of proteostatic stress.

Overall, our findings establish RECS1 as a previously unrecognized component of the exosome biogenesis pathway. RECS1 engages with Syntenin-1 and enhances exosome production, revealing a novel function for this lysosomal membrane protein and bringing to light new avenues for understanding how cells coordinate degradative and secretory pathways under stress.

## MATERIAL AND METHODS

### Antibodies and reagents

| ANTIBODY (technique) | SOURCE | IDENTIFIER |
| --- | --- | --- |
| Anti-GAPDH (WB) | Santacruz | sc-365062 |
| Anti-Flag (WB, IF) | Sigma Aldrich | F1804 |
| Anti-TMBIM1, mouse (WB, IF) | MyBioSource | MBS1499661 |
| Anti-TMBIM1, rabbit (PLA) | Abcam | ab121358 |
| Anti-Syntenin-1 (WB, PLA) | Abcam | ab19903 |
| Anti-Calnexin (WB) | Enzo Biosciences | ADISPA865 |
| Anti-HA (WB) | Santacruz | SC-7392 |
| Anti-myc (WB) | Abcam | ab9106 |
| REAGENTS | SOURCE | IDENTIFIER |
| Doxycycline | Sigma Aldrich | D9891 |
| Blasticidin | Sigma Aldrich | 203350 |
| Puromycin | Sigma Aldrich | P9620 |
| G418 | Sigma Aldrich | G8168 |
| Chloroquine | Sigma Aldrich | C6628 |
| Flag-magnetic beads | Thermo Fisher | A36797 |
| HA-magnetic beads | Thermo Fisher | 88837 |
| myc-magnetic beads | Thermo Fisher | 88842 |
| Fluoromount | Sigma Aldrich | F4680 |
| Hoechst 34580 | Thermo Fisher | H21486 |
| Propidium iodide | Invitrogen | P3566 |
| Effectene transfection reagent | Qiagen | 301425 |
| Dulbecco's modified Eagle's medium (DMEM) | Thermo Fisher | 10564011 |
| Fetal bovine serum (FBS) | Thermo Fisher | A5669701 |
| Non-essential amino acids | Thermo Fisher | 11140050 |
| Paraformaldehyde | Merck | 818715 |
| Triton X-100 | Merck | 58603 |
| Bovine serum albumin (BSA) | Merck | 12659-500GM |
| Tween-20 | Sigma Aldrich | 8.221.841.000 |
| Protease inhibitor cocktail | Sigma Aldrich | 4693124001 |
| Triethanolamine | Merck | T58300 |
| Dimethylpimelinediimidate | Merck | 80490 |
| Duolink® In Situ Red Starter Kit | Sigma Aldrich | DUO92101 |

### Cell culture and compounds

All MEF, HeLa, and human embryonic kidney (HEK) cells were maintained in Dulbecco’s modified Eagle’s medium supplemented with 10% fetal bovine serum and nonessential amino acids and grown at 37°C and 5% CO_2_.

### Transfections and DNA constructs

Syntenin-1-HA and RECS1-myc plasmids were transiently transfected with Effectene following the manufacturer’s protocol. Cell processing and treatments were performed after 48 h of transfection. Cells bearing the Flag-RECS1 doxycycline-inducible plasmid system were generated previously^1^.

### Western blot

Cells were collected and homogenized in radioimmunoprecipitation assay buffer. Total cell protein extracts were quantified and assessed by western blot as described^1^. The antibodies were diluted in blocking solution [phosphate-buffered saline (PBS) and 0.1% Tween 20 containing 5% milk]. Bound antibodies were detected using peroxidase-coupled secondary antibodies and the enhanced chemiluminescence system.

### Immunocytochemistry

Cells grown in coverslips were fixed with 4% paraformaldehyde at RT for 10 m, washed twice with PBS, permeabilised with 0.5% Triton X-100 at RT for 10 m, washed twice with PBS, blocked with 5% BSA at room temperature (RT) for 1 h, washed twice with PBS, incubated with primary antibodies diluted 1:400 in 0.5% BSA at RT for 1 h, washed twice with PBS, incubated with secondary antibodies diluted 1:1000 in 0.5% BSA at RT for 1 h, washed twice with PBS, incubated with Hoechst diluted 1:10,000 in 0.5% BSA at RT for 5 minutes, washed twice with PBS and once with distilled water and mounted on glass slides with Fluoromount before taken to a Nikon C2+ Eclipse Ti confocal microscope.

### Immunoprecipitation

After treatments and/or stimulations, cells were washed twice with PBS, collected by trypsinization and lysed in buffer containing 1% CHAPS, 20 mM Tris HCl (pH 8), 150 mM NaCl, and protease inhibitor cocktail. Magnetic beads conjugated with Flag or Myc were equilibrated with two washes of 0.2 M triethanolamine (pH 8.2), crosslinked with 20 mM dimethylpimelinediimidate diluted in 0.2 M triethanolamine (pH 8.2) for 30 minutes in shaker at room temperature and incubated with 50 mM tris (pH 7.5) in shaker for 15 minutes at room temperature. Beads were washed three times with 0.05% PBS-Tween-20 and incubated with lysates (2000 µg for mass-spectrometry experiments and 500 µg for western blot experiments) for 16 h at 4°C under rotation. Then, beads were washed thrice with 0.05% PBS-Tween-20, and proteins were eluted from beads with 2x LDS for 10 minutes at 55°C on a shaker.

### Interactome by IP-MS

Interactome analysis was performed using an immunoprecipitation-based pulldown followed by mass spectrometry on a ZenoTOF 7600 system (SCIEX). A total of 2 mg of protein lysate was subjected to immunoprecipitation, and the resulting complexes were resolved by SDS-PAGE. Gel bands were excised, subjected to in-gel trypsin digestion, and the peptides were analyzed on the ZenoTOF 7600 mass spectrometer. Raw data were processed using Spectronaut 17 (Biognosys) and searched against the pan-human library provided with the software. Proteins were considered significantly enriched if they met the thresholds of log₂(fold change) ≥ 3 and q-value ≤ 0.001.

### Cell death analysis

Extent of cell death in cells following different treatments was analysed by FACS following staining of the cells with propidium iodide. Cells were lifted from plates along with their culture media and 5 µL of propidium iodide diluted 1:1000 was added prior to assessment on a BD FACSCanto II.

### Proximity ligation assay

Proximity ligation assay (PLA) was performed using the Duolink® In Situ Red Starter Kit according to the manufacturer’s instructions, with minor modifications. Briefly, cells were fixed with 4% paraformaldehyde for 15 minutes at room temperature, permeabilised with 0.1% Triton X-100 for 10 minutes, and blocked in Duolink® blocking solution for 1 h at 37°C. Samples were incubated with primary antibodies raised in different species overnight at 4°C. After washing, species-specific PLA probes (PLUS and MINUS) were applied for 1 hour at 37°C, followed by ligation and amplification reactions. PLA signals were visualized as distinct fluorescent puncta using a Nikon C2+ Eclipse Ti confocal microscope with the pinhole altered to 90 μm. Nuclei were counterstained with DAPI, and images were analysed using ImageJ. Negative controls omitting one or both primary antibodies were included to confirm specificity.

### Extracellular vesicles and exosome purification

Four 15 cm cell culture dishes were cultivated until reaching approximately 75% confluency. Cells were exposed to DMSO or doxycycline for 16 h. The next day, dishes were washed twice with PBS and media was replaced with DMEM without FBS for 24 h, doxycycline was added back. The following day media was collected and processed as described^2^. The whole cell lysate and microvesicles fractions were frozen for posterior analysis and the exosome-containing media was kept at 4°C until ultracentrifugation. Samples were centrifuged at 100,000 g for 4 h. Supernatant was removed, and the pellet was resuspended in filtered PBS, which was then ultracentrifuged at 100,000 g for 2 h on a Thermo Scientific WX100+ ultracentrifuge with a T-1250 rotor for 30 mL tubes. Supernatant was removed, and the pellet was resuspended in 100 μL of PBS, for NTA analysis or in RIPA buffer for immunoblot.

### Nanosight exosome quantification

Ultracentrifuged exosomal fractions resuspended in 100 μLof filtered PBS were diluted 1:5. The 500 μL were passed through a NanoSight NS300. 3 videos of 30 s were captured and analysed for particle size and number on NanoSight NTA software. The number of particles were normalized to the protein concentration of whole cell lysate from the originating cells.

### Fly stocks

The following stocks were used for AGs experiments: y^1^, w*; NP5130-Gal4 (B93857); w*; UAS-EGFP::CD63; Dr^1^/TM3, Sb^1^ (B91390); y-IR/CyO (B64527); y^1^, v^1^; UAS-dRecs1 TRiP/TM3, Sb^1^ (B28052); y^1^, w*; UAS-dRecs1-Flag (from Glavic’s laboratory)^3^. All crosses were performed at 25°C following a 12-h light/dark cycle.

### Immunofluorescence of Accessory Glands (AGs)

Immunofluorescence of accessory glands was performed following the protocol described by Corrigan et al. 2014, with the exception that the animals analyzed did not express temperature-sensitive GAL8[ts]^23^. Briefly, males were selected two days post-hatching and mated to virgin females overnight. Females were then removed and left alone for one day before being dissected. Accessory glands were fixed in 4% PFA for 15 minutes, and the protocol described by Corrigan et al. was followed. To label the septate junctions, the primary antibody mouse anti-Fas3 (1:10; Developmental Studies Hybridoma Bank) was used, then an Alexa Fluor 633 antibody (1:250) as a secondary antibody. CD63::GFP levels were analyzed by direct fluorescence confocal microscopy, while nuclei were stained with DAPI (1:100000). The z-stacks were captured using an LSM 710 confocal microscope.

### Quantification of extracellular vesicles (EVs)

Image stacks (voxel size: 0.346 µm lateral, 4 µm axial) were analyzed in Imaris using batch processing. The Fasciclin 3 (red) channel was median filtered to enhance gland boundaries, then used to segment the accessory gland as a 3D surface (15 µm smoothing); the resulting surface was manually adjusted for accuracy. CD63::EGFP-labeled intraluminal vesicles (ILVs) and MVBs were segmented as secondary surfaces (detail level 0.5 µm; minimum separation 1.5 µm). EVs were detected with the Imaris Spots tool (estimated diameter 0.2 µm), with masking applied to exclude any spots inside ILVs and MVBs (ensuring only EVs outside MVBs were counted). For each larva (n = 7), gland surface area, total MVB surface area, and EV count were measured (EV counts were normalized to gland area). 3D renderings of each sample were generated to visualize the gland surface, MVBs, and EV spots.

### Bioinformatic analysis

STRING analysis was performed with STRING version 10 (http://version10.string-db.org/) using the hits that had an average log2 ratio of the two compared conditions higher than 3.00, with a k-means clustering of 3. Gene ontology analysis for cellular components was performed with a maximum false discovery rate (FDR) ≤ 0.05 and a minimum strength ≥ 0.01.

### Statistics

Results were statistically analysed an using unpaired t-test. In all plots, P values are shown as *P < 0.05, **P < 0.01, ***P < 0.001 were considered significant. In vitro data are presented as the means ± SEM. Analyses were performed using PRISM software. *D. melanogaster* data were analysed by one-way ANOVA after verifying that the data followed a normal distribution.

## ACKNOWLEDGEMENTS

We thank Dr Fedor Berditchevski (University of Birmingham, UK) for kindly providing the Syntenin-1 plasmids.

## FUNDING

FONDECYT-ANID postdoctoral fellowship # 3200718 (MM)

Millennium Institute CGR ICN2021-044, ANID (EM)

Millennium Institute CGR ICN2021-044, ANID (AG)

We acknowledge the generous support from SCIEX for the ZenoTOF 7600 mass spectrometer at the Buck Institute (BS)

NIH grants R01AG061879 and P01AG066591(LME)

U.S. Air Force Office of Scientific Research FA9550-21-1-0096 (CH)

FONDAP program 15150012

FONDECYT [1220573]

ECOS-ANID [ECOS230024]

US Army Medical Research Acquisition Activity (USAMRAA) project number AL2201415

DoD Award HT9425-23-1-0990, AL220141

## AUTHOR CONTRIBUTIONS

Conceptualization: MM, CH

Experimentation: MM, NM, CS, JB, EM

Interpretation: MM, CH, JB, EM, AG, BS, LME

Supervision: CH, AG, BS, LME

Writing – original draft: MM

Writing – review & editing: MM, CH, AG, BS, LME

## COMPETING INTERESTS

All authors declare they have no competing interests.

## DATA AND MATERIAL AVAILABILITY

All data needed to evaluate the conclusions in the paper are present in the paper and/or the Supplementary Materials. Information and requests about reagents and resources should be directed to the lead contact, Claudio Hetz.

---

**Supplemental Figure 1.**
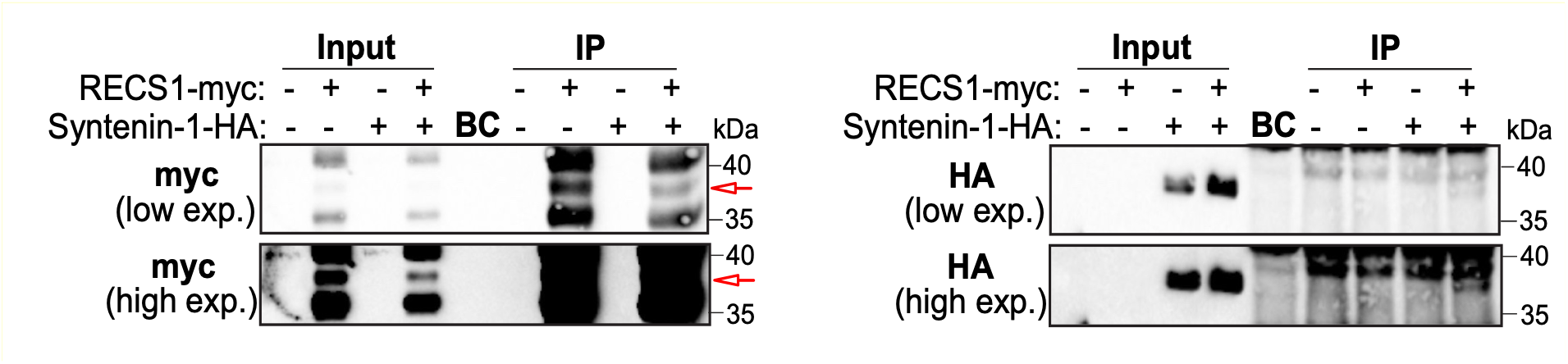
HEK-293 cells transfected with RECS1-myc and/or Syntenin-1-HA plasmids were processed for HA immunoprecipitation. The red arrow on the left panel points to the RECS1-myc band. Syntenin-1-HA on the right panel is found by the 35 kDa marker.

**Supplemental Figure 2.**
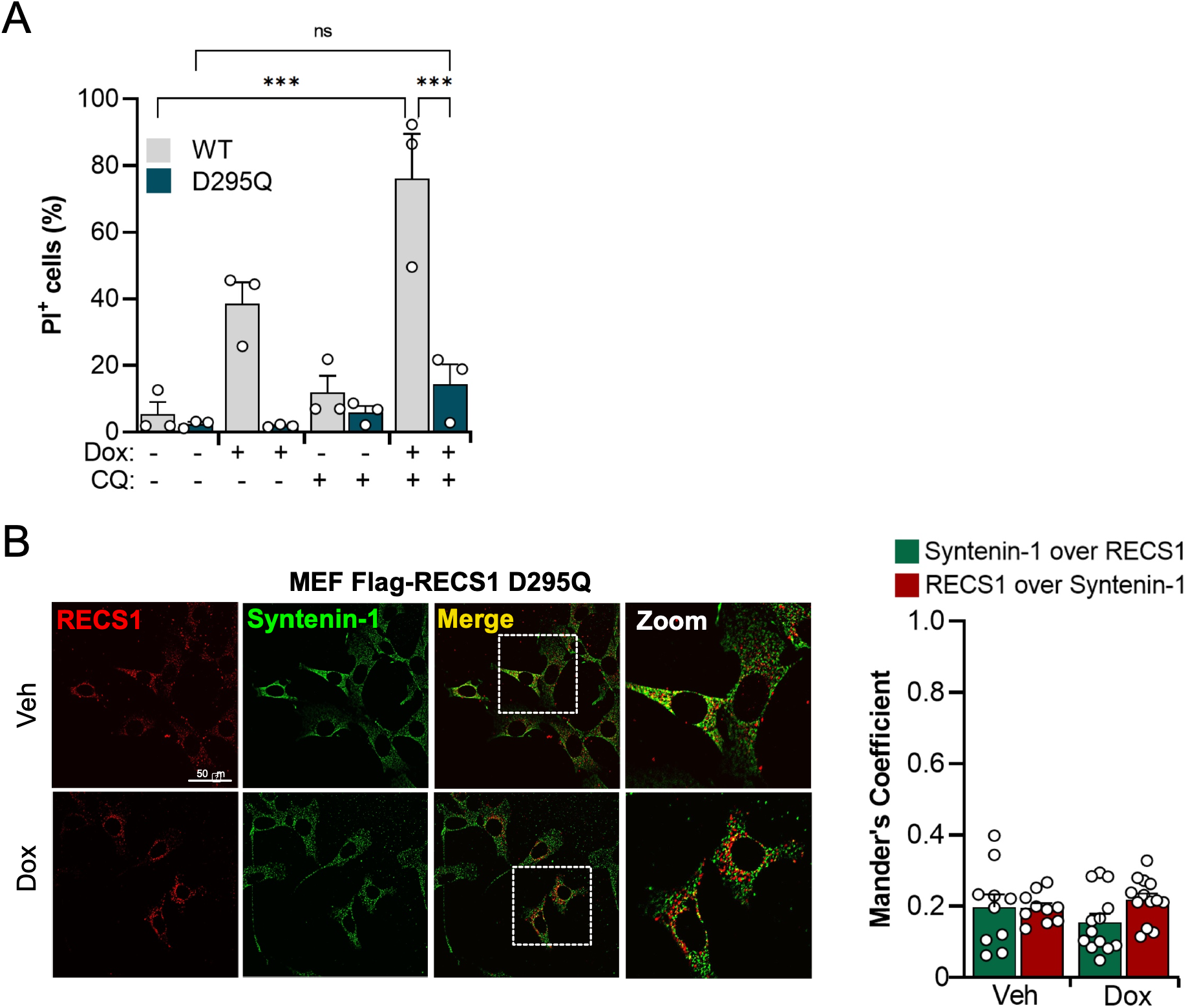
(A) MEF Flag-RECS1 and Flag-RECS1 D295Q cells were stimulated with doxycycline (1 µg/ml, 16 h), chloroquine (50 µM, 24 h), or a combination of both and analysed for apoptosis by PI staining on FACS. **(B)** MEF Flag-RECS1 D295Q were stimulated with doxycycline (1 µg/ml, 16 h), processed for immunocytochemistry and stained for RECS1 and Syntenin-1. Left: representative confocal images. Right: co-localization quantification by Mander’s coefficient; green depicts the percentage of signal of Syntenin-1 over RECS1 and red depicts the percentage of signal of RECS1 over Syntenin-1.

## Notes

### Competing Interest Statement

The authors have declared no competing interest.

